# Analysis of histone H3K4me3 modifications in bovine placenta derived from different calf-production methods

**DOI:** 10.1101/2023.07.01.547357

**Authors:** Kanoko Hattori, Yoichiro Hoshino, Masayuki Kachi, Yasumitsu Masuda, Satoshi Yamamoto, Shinnosuke Honda, Naojiro Minami, Shuntaro Ikeda

## Abstract

In ruminants, overgrowth of offspring produced by in vitro fertilization (IVF) is a common problem. Abnormal epigenetic modifications caused by environmental factors during the early embryonic period are suspected as an etiology of overgrowth. In this study, we investigated the genome-wide histone H3K4me3 profiles of bovine placentae that play a pivotal role in fetal development and compared their characteristics between artificial insemination (AI)- and IVF-derived samples. Cotyledons were harvested from the placentae obtained at parturition of 5 AI- and 13 IVF-derived calves, and chromatin immunoprecipitation sequencing was performed for H3K4me3. We confirmed no significant maternal tissue contamination in the samples we used. The revealed H3K4me3 profiles reflected the general characteristics of the H3K4me3 modification, which is abundantly distributed in the promoter region of active genes. By extracting common modifications from multiple samples, the genes involved in placenta-specific biological processes could be enriched. Comparison with the H3K4me3 modifications of blastocyst samples was also effective for enriching the placenta-specific features. Principal component analysis suggested the presence of differential H3K4me3 modifications in AI- and IVF-derived samples. The genes contributing to the difference were related to the developmental biological processes. Imprinted genes such as *BEGAIN, ZNF215*, and *DLX5* were among the extracted genes. Principal component and discriminant analyses using only male samples categorized the samples into three groups based on fetal weight and calf-production methods. To our knowledge, this is the first study to profile the genome-wide histone modifications of bovine fetal placentae and reveal their differential characteristics between different calf-production methods.

## Introduction

The prevalence of assisted reproductive technologies (ARTs), both in the field of human medicine and in the production of resource animals, has been remarkable, enabling a wide variety of ways for humans and animals to be born. Although the benefits of ART are immeasurable, we need to be aware of developmental anomalies attributed to ART. For example, in ruminant livestock production, fetal overgrowth, also known as large offspring syndrome (LOS), is a more frequent problem in offspring derived from in vitro fertilization (IVF) than in those derived from artificial insemination (AI), where fertilization occurs in vivo (Chen et al., 2015; van Wagtendonk-de Leeuw, Aerts, & den Daas, 1998; Young, Sinclair, & Wilmut, 1998). The etiology of LOS is not clearly understood, but abnormal epigenetic modifications caused by environmental factors during early embryonic period are suspected (Chen et al., 2015; Chen, Hagen, Ji, Elsik, & Rivera, 2017). In particular, the misregulation of imprinted genes such as *IGF2R* and *KCNQ1OT1* is widely observed in tissues of overgrown fetuses born from IVF (Chen et al., 2015). However, most such misregulation of gene expression is not explained by changes in DNA methylation (Chen et al., 2017), and thus it is possible that IVF may cause alterations in epigenetic modifications other than DNA methylation, leading to the misregulation of gene expression.

As the interface between mother and fetus, the placenta plays a pivotal role in fetal development (Burton & Fowden, 2015). Because part of the placenta is derived from the fetal lineage (Thowfeequ & Srinivas, 2022), it may epigenetically inherit the effects of environmental differences during the embryonic period caused by ART interventions. The structure of the placenta differs among species, but in ruminants such as cattle, the placenta has a polyplacental structure with multiple placentae scattered over the placental membrane that surrounds the fetus. The villous plexus (cotyledon) of the placenta on the fetal side is in close contact with a vascular-rich ridge (caruncle) on the maternal uterine epithelium. This interdigitated unit is called the placentome (Carlson et al., 2021). The cotyledon is rich in blood vessels, through which substances are exchanged with the mother. In particular, it has been shown that placental villi and capillaries are more developed in the second half of gestation, when the need for nutritional supply through the placenta increases (Leiser et al., 1997). This indicates that the cotyledon is an important tissue for maintaining the nutritional status of the fetus. Furthermore, placental tissue is an extra-embryonic tissue that can be collected in a non-invasive manner both for the fetus and the mother because it is normally delivered spontaneously from the mother at parturition. Genome-wide epigenomic analyses are underway in livestock animal tissues (Han, Ren, Cao, Zhao, & Yu, 2019; Kern et al., 2021; Villar et al., 2015; Zhao et al., 2015), but there are no data on bovine placental tissues. In this study, we collected the genome-wide profiles of representative permissive histone epigenetic modifications, H3K4me3, in cotyledons from term placentae and compared their characteristics between AI- and IVF-derived calves.

## Materials and Methods

### Placenta collection

This study was approved by the Animal Research Committee of Kyoto University (permit numbers R3-103, R4-103, and R5-103) and was carried out in accordance with the Regulation on Animal Experimentation at Kyoto University.

AI was performed to impregnate 5 Japanese Black cows in accordance with their natural estrus cycle. Pregnancy was also induced in 13 cows by the transfer of IVF-derived Japanese Black cow embryos produced by ovum-pick-up. Information on the resulting calves is shown in Table 1. After parturition, the delivered secundinae were retrieved and transported to the laboratory while being cooled. The cotyledons were cut into small pieces, snap-frozen, and stored at −80°C prior to chromatin extraction.

**Table 1.** The list of cotyledons analyzed

| ID | Calving method* | Sex | Calf birth weight |  |  |
| --- | --- | --- | --- | --- | --- |
|  |  |  | (kg) | Stillbirth | Overweight** |
| Male 1 | AI | male | 35.9 |  |  |
| Male 2 | AI | male | 27.8 |  |  |
| Male 3 | AI | male | 31.9 |  |  |
| Male 4 | IVF | male | 50 |  | + |
| Male 5 | IVF | male | 50 |  | + |
| Male 6 | IVF | male | 52.5 | + | + |
| Male 7 | IVF | male | 60 | + | + |
| Male 8 | IVF | male | 31.1 |  |  |
| Male 9 | IVF | male | 36 |  |  |
| Male 10 | IVF | male | 37.3 | + |  |
| Female 1 | AI | female | 31 |  |  |
| Female 2 | AI | female | 24.4 |  |  |
| Female 3 | IVF | female | 37 |  | + |
| Female 4 | IVF | female | 45 |  | + |
| Female 5 | IVF | female | 39 |  | + |
| Female 6 | IVF | female | 36.7 |  | + |
| Female 7 | IVF | female | 43.5 |  | + |
| Female 8 | IVF | female | 42 |  | + |

\*AI, artificial insemination; IVF, in vitro fertilization
\*\*Heavier than the upper limits in Japanese Feeding Standard of Beef Cattle (2008)

### ChIP-seq

ChIP was performed using a SimpleChIP Plus Sonication Chromatin IP Kit (Cell Signaling Technology, Danvers, MA) according to the manufacturer’s protocol with some modifications. Approximately 200-mg samples (cotyledons) were crosslinked with 1% formaldehyde for 10 min at room temperature and the crosslinking was quenched by glycine. The samples were homogenized using BioMasher (Nippi, Tokyo, Japan) in 1x ChIP Sonication Cell Lysis Buffer. After centrifugation and supernatant removal, the samples were soaked in ChIP Sonication Nuclear Lysis Buffer and sonicated to shear chromatin using a Bioruptor UCD-250 (Cosmo Bio) for 30 cycles of 30 s each with 30-s pauses in ice water. The sample was centrifuged for 10 min at 21,000 × g and the supernatant was transferred to a new tube. Approximately 10 μg chromatin was aliquoted and diluted to 500 μL of 1x ChIP Buffer and immunoprecipitated using 1.4 μg anti-H3K4me3 antibody (pAb-003-050, Diagenode, Denville, NJ). In some cases, 200 ng of chromatin was set aside as “input.” The input and ChIPed DNA were de-crosslinked, purified, and eluted in 50-μL DNA Elution Buffer. The DNA (40–50 ng) samples were processed for library preparation for next-generation sequencing by using a Next Gen DNA Library Kit and Next Gen Indexing Kit (Active Motif, Carlsbad, CA) following the manufacturer’s protocol.

Sequencing was performed on a HiSeqX (Illumina, San Diego, CA) as paired-end 150-base reads. The sequencing reads were quality-checked, merged, and aligned to the bovine genome (Bos_taurus_UMD_3.1.1/bosTau8, June 2014) using Bowtie 2. Handling of sam and bam files was performed using Samtools (http://www.htslib.org/). Mapping duplicates were removed by Picard (http://broadinstitute.github.io/picard/). The generated bam files were converted to bigWig (bw) files by the bamCoverage tool of deepTools (https://deeptools.readthedocs.io/en/develop/) with counts-per-million normalization. The H3K4me3 landscapes were visualized using Integrative Genomics Viewer (Robinson et al., 2011). The H3K4me3 peaks were called using MACS2 (Liu, 2014; Zhang et al., 2008). The annotation of called peaks to genomic regions was generated using CEAS (Shin, Liu, Manrai, & Liu, 2009). Average and gene-specific H3K4me3 signal profiles were generated using ngs.plot (Shen, Shao, Liu, & Nestler, 2014) and the generated data were used to compare H3K4me3 levels at specific genes after normalization such that the height of the highest point of the average profile was set to 1.

### RNA-seq

To evaluate the gene expression levels in some samples, RNA-seq was performed as follows. Total RNA was isolated using RNeasy Fibrous Tissue Mini Kit (Qiagen, Hilden, Germany) according to the manufacturer’s protocol. Approximately 20 mg cotyledons were homogenized using BioMasher. Removal of rRNA as well as mitochondrial RNA was performed using RiboGone™ - Mammalian (Takara Bio USA, Inc., San Jose, CA) from total RNA according to the manufacturer’s protocol. The ribosomal RNA-depleted RNA (5–10 ng) was used to generate sequencing libraries, using SMARTer^®^ Stranded RNA-Seq Kit (Takara Bio USA, Inc.) following the manufacturer’s protocol. The first-strand cDNA was purified using SPRIselect beads (Beckman Coulter, Brea, CA) and purified cDNA was amplified into RNA-seq libraries by PCR with 15 cycles, regardless of the input quantity. Then the amplified RNA-seq library was purified using SPRIselect beads again and eluted in 20 μL Stranded Elution Buffer. Sequencing was performed in the same manner as ChIP-seq, as described above. The sequencing reads were quality-checked, trimmed, and aligned to the bovine genome (Bos_taurus_UMD_3.1.1/bosTau8, June 2014) using HISAT2 (Kim, Langmead, & Salzberg, 2015; Kim, Paggi, Park, Bennett, & Salzberg, 2019). Handling of sam and bam files as well as generating bigwig (bw) files were performed in the same manner as ChIP-seq, as described above. Quantification of gene expression as FPKM was performed by String-Tie (Pertea et al., 2015).

### Data analyses

The DAVID tool (Huang da, Sherman, & Lempicki, 2009a, 2009b) was used for gene ontology analysis. For normalized ChIP-seq signals around the transcription start site (TSS) ± 1000 bp, principal component analysis (PCA) and partial least squares-discriminant analysis (PLS-DA) were performed using MetaboAnalyst 5.0 (Pang et al., 2021; Pang et al., 2022) after narrowing the number of genes to those with a coefficient of variation in the top 5,000. Data were analyzed after filtering based on the interquantile range (default) and subsequent auto-scaling.

## Results

### Assessment of maternal genomic contamination

To ensure that the genome-wide H3K4me3 data obtained in this study were derived from tissue on the calf side of the placenta, the possibility and extent of maternal tissue contamination were examined as follows. First, we focused on the overall signals of the H3K4me3 in autosomes and the X chromosome (Fig. 1a). The 10 male samples exhibited H3K4me3 signals on the X chromosomes that were approximately half of those on the 29 autosomes and the female X chromosome, suggesting that these 10 samples had only one X chromosome. However, the amount of signals on the autosomes in the female samples were nearly identical to those on the X chromosomes. These results reflect the fact that bovine sex chromosomes are of the male heterozygous type, as in other mammals. Next, we turned our attention to *XIST*, which is transcribed from one of the two X chromosomes in females and inactivates the chromosome from which *XIST* transcription occurs (Markaki et al., 2021). Although clear H3K4me3 modification peaks were observed near the transcription start site of *XIST* in placental samples from female calves, they were not observed in those from male calves (Fig. 1b). These results may reflect female-specific X chromosome inactivation. Furthermore, H3K4me3 modification on the *PRAME* gene was observed in placental samples only from male calves (Fig. 1c). *PRAME* is an ancestral gene of the *Oog* family observed in mice, and it is known that *PRAME* on chromosome 17 in cattle has been translocated and amplified on the Y chromosome (Chang et al., 2011). The absence of an H3K4me3 modification signal near *PRAME* in placental samples from female calves also enabled clear differentiation between male and female tissues, suggesting that there was no contamination of maternal *PRAME* sequence in the male samples.

**Fig. 1.**
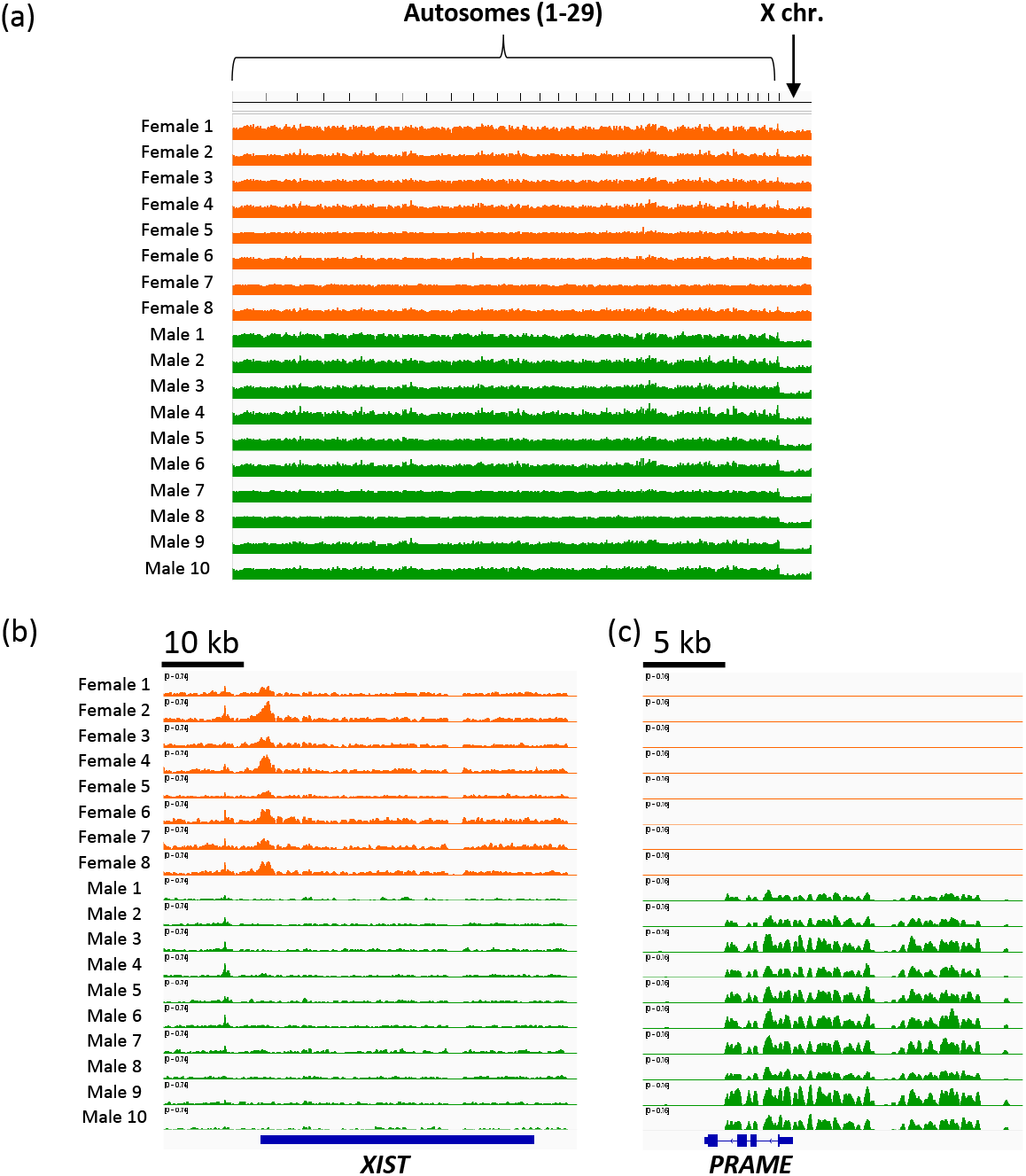
Verifying that there was no significant contamination of the maternal genome in the ChIP-seq analysis. (a) Overall signals of the H3K4me3 in autosomes and the X chromosome. (b) H3K4me3 signals at the *XIST* gene. (c) H3K4me3 signals at the *PRAME* gene.

### Overview of genome-wide H3K4me3 profiles of bovine placentae

We performed ChIP-seq analysis of H3K4me3 using cotyledons (n = 18, Table 1) derived from placentomes of full-term placentae collected at parturition. When the criteria for being overweight were set according to the Japanese feeding standard for beef cattle (*National Agriculture and Food Research Organization, Japanese Feeding Standard for Beef Cattle* 2008), overweight was clearly more frequent in IVF-derived calves (Fisher’s exact test, P < 0.01). Figure 2a shows a snapshot of the H3K4me3 landscape in a 92-kb region of chromosome 5 that encompasses the TSS of some genes including *GAPDH*, a representative positive region for H3K4me3 modifications. However, such enrichment did not occur in the input samples for which ChIP was not performed, indicating the specificity of the analysis. Approximately 22% of the called peaks were located at the promoter regions of the genome, demonstrating clear enrichment of the H3K4me3 modifications in these regions (Fig. 2b). Average profile plotting around the TSS categorized by gene groups with different expression levels showed that highly expressed gene groups had more extensive H3K4me3 modifications in consistent with the general association of H3K4me3 with transcriptional activation (Fig. 2c). We extracted 10,200 H3K4me3 peaks that were common across the 18 samples. From the genes that harbored these common peaks within ± 3,000 bp of the TSS, those with the top 100 peak occupancy rates were subjected to gene ontology (GO) analysis. The most significant GO term was “regulation of gene expression.” In addition, the term “circulatory system development” was also enriched as a term closely related to the placental function (Fig. 2d).

**Fig. 2.**
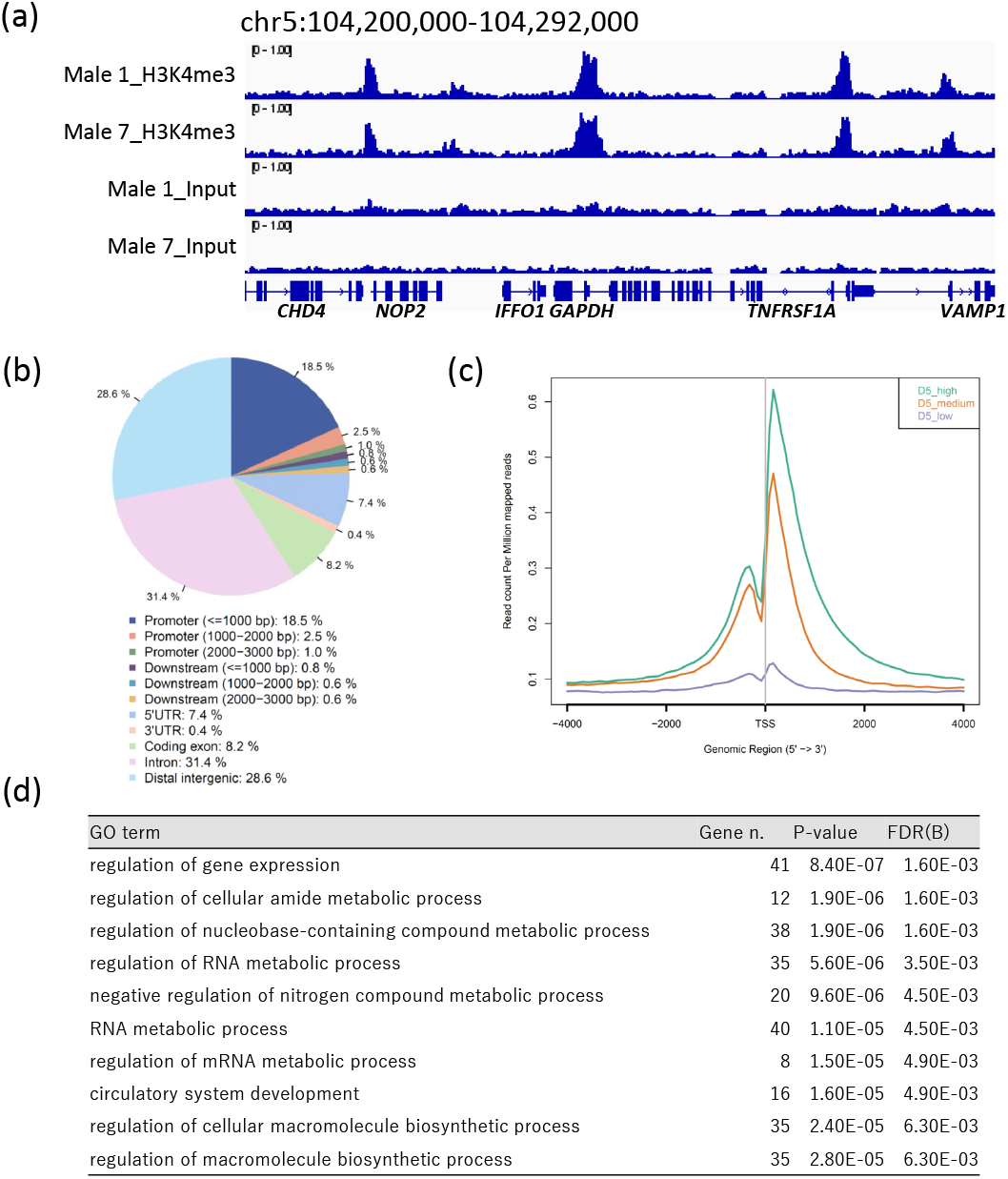
Overview of H3K4me3 ChIP-seq results in bovine cotyledons. (a) Snapshot of the H3K4me3 landscape in a 92-kb region of chromosome 5 that encompasses the transcription start site (TSS) of some genes, including *GAPDH*. The ChIP peaks in two male samples were visualized using the Integrative Genomics Viewer (Robinson et al., 2011). (b) Distribution of H3K4me3 peaks in corresponding genic and intergenic regions. The figure was generated using the CEAS tool (Shin et al., 2009). (c) Average profile plot of the H3K4me3 signal around the TSSs categorized by gene groups with different expression levels. The sample from Female 6 was used to generate the figure. The average profile plots were generated using ngs.plot (Shen et al., 2014). (d) Top 10 significant GO terms for biological processes enriched by the genes that harbored the common peaks with the top 100 highest peak occupancy rates. Gene n. represents the numbers of related genes. FDR(B) indicates the Benjamin false discovery rate.

### Characterization of placenta- and blastocyst-specific H3K4me3 modification

Using the ChIP-seq data from one placenta (Male 1), we attempted to extract H3K4me3 modifications that are specific or common against blastocysts. First, 30,257 peaks obtained from the ChIP-seq data of placenta (Male 1) were designated as placental peaks. The 13,110 common peaks from two single blastocysts (P14 and P15) previously reported from our laboratory (Susami, Ikeda, Hoshino, Honda, & Minami, 2022) were designated as the blastocyst peaks. By comparing the placental and blastocyst peaks, we extracted 18,099 placenta-specific peaks, 12,168 common peaks, and 2,921 blastocyst-specific peaks (Fig. 3a). GO analysis of these peaks revealed that the genes with placenta-specific peaks enriched the terms closely related to placental functions, including “circulatory system development” and “vasculature development.” These terms closely related to placental tissue function were not enriched in the GO analysis using placental peaks, but rather only the more general terms were enriched (Fig. 3b). This result is consistent with the results of a previous study showing that data on histone modifications in blastocysts are useful as a “sieve” through which to extract tissue features (Ishibashi, Ikeda, & Minami, 2021).

**Fig. 3.**
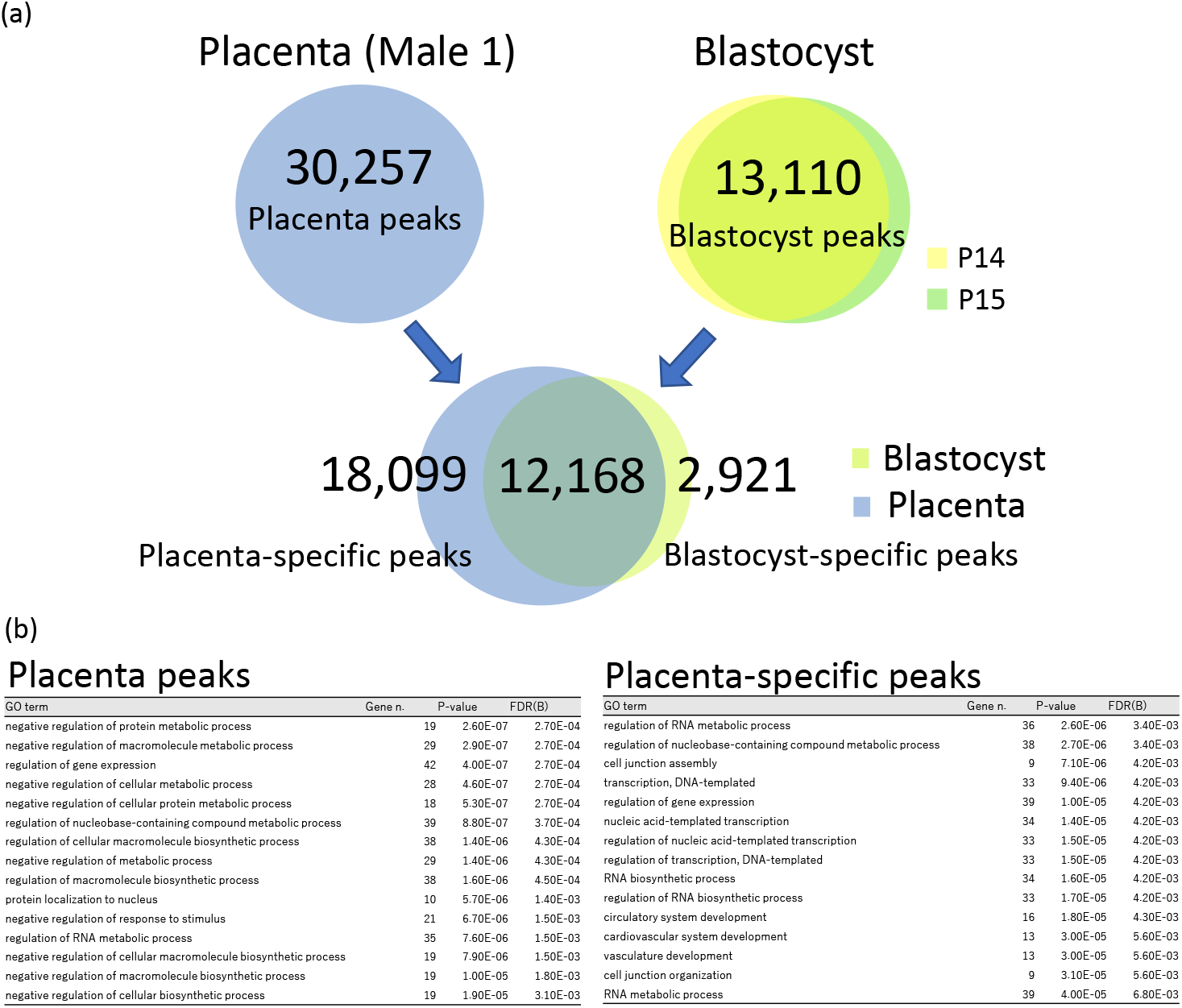
Characterization of placenta- and blastocyst-specific H3K4me3 modification. (a) Processing of ChIP-peaks in placenta and blastocyst samples. The numbers show those of the peaks. (b) Top 15 significant GO terms for biological processes enriched by the genes with the top 100 highest peak occupancy rates. Gene n. represents the numbers of related genes. FDR(B) indicates the Benjamin false discovery rate.

### Difference in H3K4me3 between AI- and IVF-derived placentae

Fig. 4a and b shows the results of PCA and PLS-DA of the normalized H3K4me3 levels at gene promoters in all the samples. The results indicated that the samples were roughly classified between AI- and IVF-derived placentae. We extracted the top 100 genes of PLS-DA VIP scores between the two groups by using heatmap analysis (Fig. 4b). GO analysis of these genes revealed enrichment of GO terms related to organismal development, including “epithelial cell proliferation,” “animal organ morphogenesis,” and “cell proliferation” (Fig. 4c). Among these 100 genes were the imprinting genes *BEGAIN, ZNF215*, and *DLX5*, which have been shown to be imprinted in cattle or in at least one other mammalian species (Fig. 4d).

**Fig. 4.**
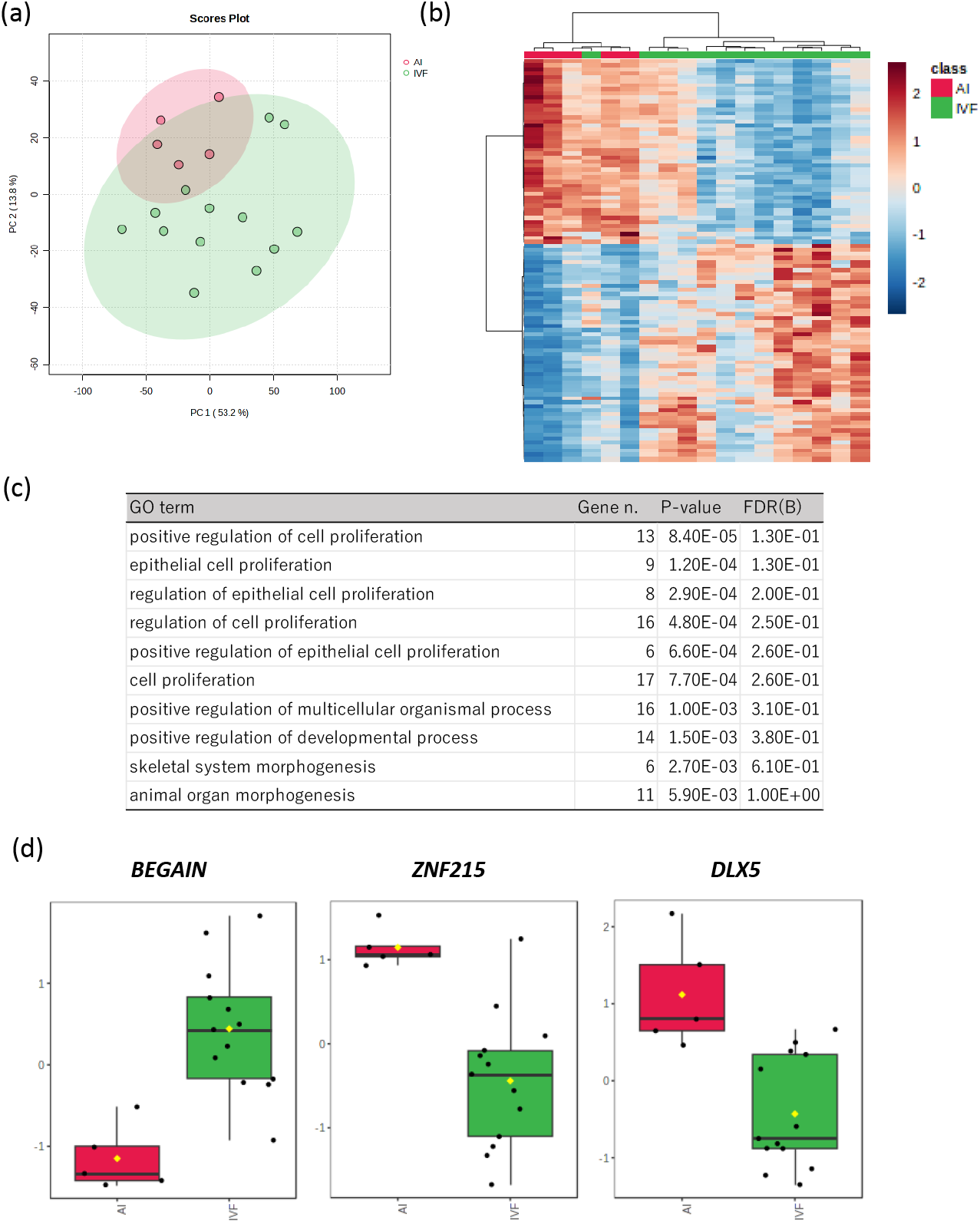
Characterization of H3K4me3 profiles between AI- and IVF-derived cotyledons. (a) Principal component analysis of 18 cotyledon samples considering the normalized peak areas around the TSSs. The pink and green shades represent the respective 95% confidence intervals. (b) Heatmap derived from top 100 genes of partial least squares-discriminant analysis (PLS-DA) VIP scores in terms of H3K4me3 around TSSs between AI- and IVF-derived cotyledons. (c) Top 10 significant GO terms for biological processes enriched by the genes indicated in (b). Gene n. represents the numbers of related genes. FDR(B) indicates the Benjamin false discovery rate. (d) Normalized levels of H3K4me3 modifications around TSSs in the *BEGAIN, ZNF215*, and *DLX5* genes.

Due to the limited sample size of females as well as consideration for making conditions identical, we performed PCA and PLS-DA using only male samples, considering “normal” or “overweight” as an additional variable. In both analyses, the three groups (AI-normal weight, IVF-normal weight, and IVF-overweight) were classified into three clusters (Fig. 5a). Figure 5b shows the heatmap analysis using the top 50 PLS-DA VIP scores that showed different levels of H3K4me3 among the groups. Representative genes for distinguishing each group included *ZC3H7B, MAP3K9*, and *NPHP1* (Fig. 5c).

**Fig. 5.**
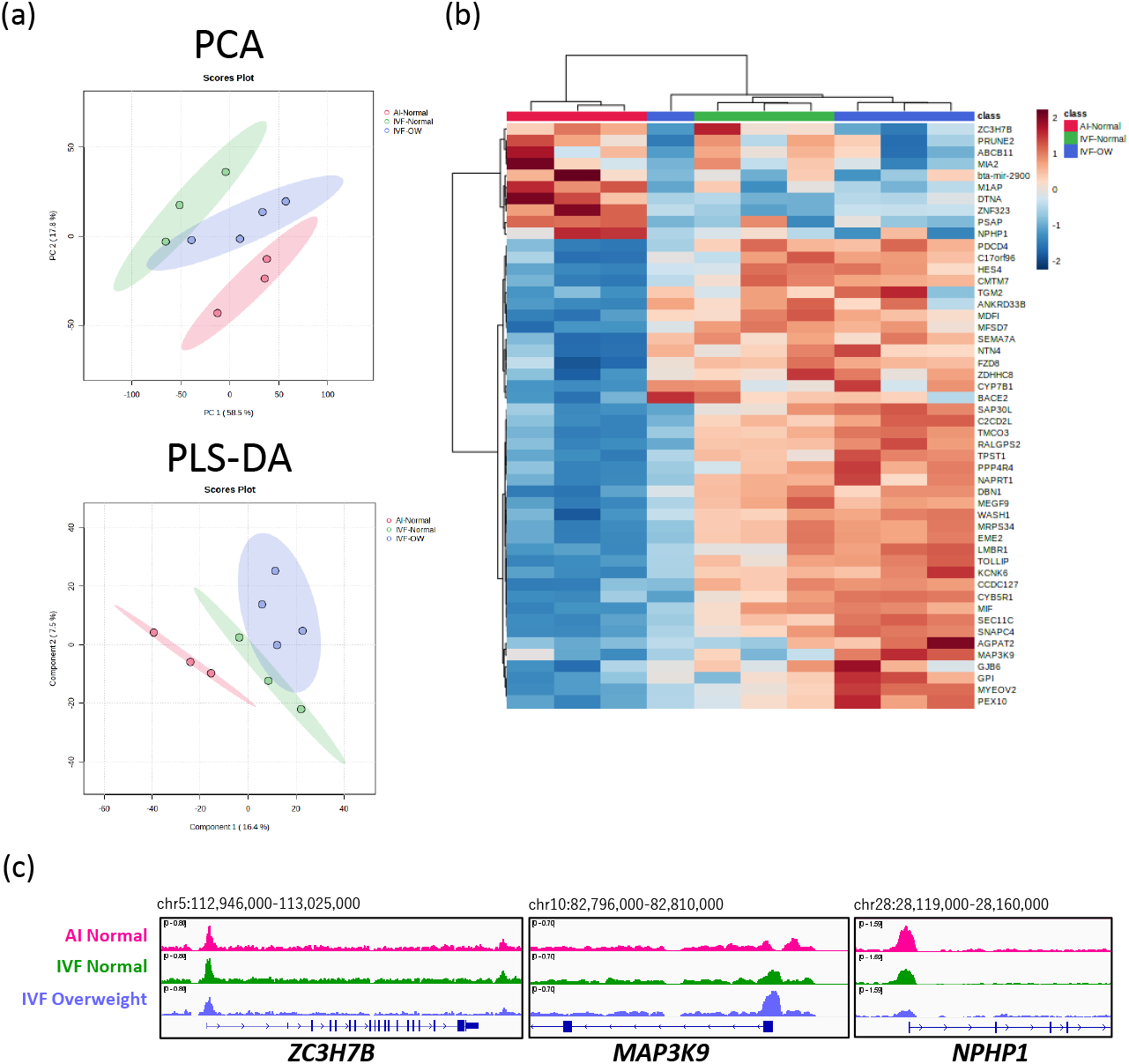
(a) Principal component analysis (PCA, upper) and partial least squares-discriminant analysis (PLS-DA, lower) of 10 male cotyledon samples categorized by AI origin (all normal weight), IVF origin with normal weight, and IVF origin with overweight (OW). The pink, blue, and green shades indicate the respective 95% confidence intervals. (b) Heatmap derived from top 50 genes of PLS-DA VIP scores. (c) The H3K4me3 landscapes of the *ZC3H7B, MAP3K*, and *NPHP1* genes in AI origin (male 3), IVF origin with normal weight (male 9), and IVF origin with OW (male 5).

## Discussion

To our knowledge, this is the first report of a genome-wide histone modification profile in bovine placental tissues. We were concerned about whether this experimental method would yield information on the fetal side of the placenta with little contamination of maternal DNA. Consequently, in terms of the overall amount of H3K4me3 modification of the X chromosome in comparison with the autosomes, the amount of H3K4me3 on the *XIST* gene, and the detection of the *PRAME* gene sequence, we confirmed that there was no significant maternal tissue contamination in the placental samples from male calves, and we consider that there was also no such contamination in the placental samples from female calves, which were sampled in the same manner as the males.

The overall H3K4me3 ChIP-seq results indicated typical characteristics of the modification represented by general enrichment in the active gene promoters (Liang et al., 2004), suggesting the validity of the experimental method. We previously reported that subtracting H3K4me3 modifications common to blastocysts from those in somatic tissues can highlight the characteristic biological processes of each tissue (blastocyst H3K4me3 as a “sieve” (Ishibashi et al., 2021)). Supporting this, in the present study, this “sieving” method was also effective for highlighting the placental characteristic biological processes represented by circulatory functions (Fig. 3). However, this feature was also highlighted by common peaks from multiple samples (Fig. 2d) without sieving. This result suggests that by accumulating data from multiple samples and excluding non-common and minor peaks, we were able to extract peaks that reflect the characteristics of the placental tissue.

The PCA and PLS-DA analyses against AI- and IVF-derived samples revealed the rough classification of these two groups, suggesting that there are characteristic H3K4me3 modifications that differ between AI- and IVF-derived samples. Interestingly, the genes involved in these characteristics enriched the biological processes related to cell and organ development (Fig. 4c). Because overgrown fetuses preferentially observed in IVF-derived fetuses also exhibit overdevelopment of organs, differences in the H3K4me3 modification levels of the genes involved in development may be implicated in such processes. Furthermore, the genes involved in these characteristic differences included the imprinting genes *BEGAIN, ZNF215*, and *DLX5*, which have been shown to be imprinted in cattle or in at least one other mammalian species (Fig. 3d). Misregulation of imprinting genes has been observed in tissues of IVF-derived overgrown offspring (Chen et al., 2015). Single-nucleotide polymorphisms (SNPs) within the bovine *ZNF215* gene are associated with bovine growth and body conformation traits, and the human *ZNF215* ortholog belongs to the imprinted gene cluster associated with Beckwith-Wiedemann syndrome, a disease similar to LOS in humans (Magee et al., 2010). In addition, *DLX5* was maternally expressed in porcine skeletal muscle, fat, lung, spleen, stomach, and small intestine. *DLX5* regulates the synthesis of c-aminobutyric acid (GABA), which might increase the secretion of growth hormone. Indeed, a SNP mutation in exon 1 of the *DLX5* gene in pig was highly associated with carcass composition and bone development traits (Cheng et al., 2008). Collectively, the histone modification of these genes in the placenta may be involved in the overgrowth of offspring born through IVF.

Although limited in number, we compared H3K4me3 modifications among the AI-normal weight, IVF-normal weight, and IVF-overweight groups using male samples. The multivariate analyses clearly categorized these three groups, and some genes contributing to their differences were identified (Fig. 5). However, given the small sample size and the apparent confounding between IVF-derived and overweight calves (Table 1), further studies with larger samples will be needed to identify genes associated with LOS.

In conclusion, this study reveals for the first time the genome-wide H3K4me3 modifications of bovine fetal placentae as well as the characteristic differences in modifications between different calf- production methods. Further research is needed to elucidate the causal relationship between overgrowth preferentially observed in IVF-derived fetuses as well as to reduce their occurrence.

## Acknowledgments

The authors are grateful to the staff at the Kyoto University Livestock Farm who assisted in the collection of placenta samples. This work was supported in part by the Japan Society for the Promotion of Science (19H03104 and 23H02363 to S.I.) and Livestock Promotional Subsidy from the Japan Racing Association.

## Conflict of interest

The authors declare that they have no conflict of interest.

## Author Contributions

K.H., Y.H., N.M., and S.I. conceived the experiments. M.K., Y.M., S.Y., and Y.H. performed calf production and sample collection from placentae. K.H. and S.I. performed sequence library preparation and deep sequencing. K.H., Y.H., S.H., N.M., and S.I. analyzed the results. H.K. and S.I. drafted the manuscript. All authors discussed the results and approved the manuscript.

## Notes

### Competing Interest Statement

The authors have declared no competing interest.

